# Volatiles from *Serratia marcescens, S. proteamaculans*, and *Bacillus subtilis* Inhibit Growth of *Rhizopus stolonifer* and Other Fungi

**DOI:** 10.1101/2020.09.07.286443

**Authors:** Derreck Carter-House, Joshua Chung, Skylar McDonald, Kerry Mauck, Jason E Stajich

## Abstract

The common soil bacteria *Serratia marcescens, Serratia proteamaculans*, and *Bacillus subtilis* produce small molecular weight volatile compounds that are fungistatic against multiple species, including the zygomycete mold *Rhizopus stolonifer* (Mucoromycota) and the model filamentous mold *Neurospora crassa* (Ascomycota). The compounds or the bacteria can be exploited in development of biological controls to prevent establishment of fungi on food and surfaces. Here, we quantified and identified bacteria-produced volatiles using headspace sampling and gas chromatography-mass spectrometry. We found that each bacterial species in culture has a unique volatile profile consisting of dozens of compounds. Using multivariate statistical approaches, we identified compounds in common or unique to each species. Our analysis suggested that three compounds, dimethyl trisulfide, anisole, and 2-undecanone, are characteristic of the volatiles emitted by these antagonistic bacteria. We developed bioassays for testing inhibition of each compound and found dimethyl trisulfide and anisole were the most potent. This work establishes a pipeline for translating volatile profiles of cultured bacteria into high quality candidate fungistatic compounds which may be useful in combination as antifungal control products.

**IMPORTANCE:** Bacteria may benefit by producing fungistatic volatiles that limit fungal growth providing a mechanism to exclude competitors for resources. Volatile production is potentially mediating long distance biological control and competitive interactions among microbes, but the specific bioactive compounds are poorly characterized. This work provides evidence that fungistatic compounds in complex blends can be identified using machine-learning and multivariate approaches. This is the first step in identifying pathways responsible for fungistatic volatile production in order to phenotype and select natural strains for biocontrol ability, or engineer bacteria with relevant pathways.

---

Advances in next-generation sequencing are enabling studies of how fungal and bacterial communities coexist, communicate and cooperate. Since 2018 over 300 published studies have documented communication and coexistence mechanisms of bacteria and fungi Deveau et al. (1). For example, metagenomic approaches that quantify the presence or absence of different organisms in a community are producing in-sights into the symbioses of fungi with cyanobacteria or algae partners to form lichens Meiser et al. (2). Competitive interactions and population dynamics among bacteria and fungi in the topsoil have been revealed using metagenomic approaches to characterize niche occupation and global distribution, Bahram et al. (3) and community make-up of microbes in extreme environments Gonzalez et al. (4). It should be noted that fungi are also producing volatiles and extensive work shows a multitude of compounds are produced Inamdar et al. (5). The improved understanding of cooperative, neutral, and competitive interactions among bacteria and fungi has fueled efforts to pinpoint the molecularand chemical mechanisms mediatingthese interactions. These efforts are revealing the importance of small molecules and secreted peptides as mediators enabling each interacting partner to sense, and react to, other interacting partners Schmidt et al. (6).

Beyond contributing to our fundamental understanding of microbial communities, knowledge of the molecular signals underlying bacterial-fungal interactions can directly inform efforts to control problematic fungi on food and in agricultural contexts Sathe et al. (7), Poppe et al. (8), Frey-Klett et al. (9). For example, the bacterial effectors Tfe1 and Tfe2 are produced by *Serratia marcescens* in response to the presence of *Candida albicans* (a common human fungal pathogen), reducing the fungal population by two orders of magnitude under lab conditions Trunk et al. (10). These effectors were among the first discovered to be produced in response to fungi and act to suppress fungal growth. Metabolites and enzymes produced by *Serratia* living on or near plant roots are effective at reducing pathogen populations Vleesschauwer (11). As part of the arms race between partners, having the capability to perceive, avoid, or respond to bacteria is advantageous for fungi that encounter these microbes in the soil. The fungus *Aspergillus nidulans* activates secondary metabolite pathways in response to physical contact with *Streptococcus* which may provide a defense mechanism. These metabolites are similar to lecanoric acid, which has been proposed to inhibit ATP production in bacteria. Interestingly these same metabolites are expressed in symbiosis of lichenized fungi and may serve a role in selecting compatible partners Umezawa et al. (12). The natural ability of many microbes to produce and secrete potent anti-fungal compounds has led to the commercialization of some microbes as biological control agents. For example, preparations of *Serratia plymuthica* are marketed as a strawberry disease biocontrol (Rhizostar ®, e-nema GmbH, Raisdorf, Germany) that reduces Verticillium wilt by up to 18.5% and *Phytophthora cactorum* root rot by up to 33.4%. The molecular mechanisms of fungal repression by *S. plymuthica* is not known, but may involve secondary metabolites such as prodigiosin, siderophores, haterumalides, and the production of degradation enzymes including glucanases, chitinases, and proteases Kurze et al. (13).

Bacteria-fungi interactions can be exploited for development of biocontrol strategies of fungi in medical or agricultural environments. Biocontrol can be an important resource for postharvest protection of fruits and vegetables. Filamentous fungi, in particular *Rhizopus stolonifer*, are common postharvest rots of fruits and vegetables including stone fruits, grapes, berries, tropical fruits, tomatoes, and tubers Coates and Johnson (14) and can reduce the store shelf life of many fleshy fruits and vegetables Wang et al. (15). Typical control strategies to prevent postharvest rot include storage in high CO_2_ and cold temperatures to slow proliferation of *Rhizopus* during storage and transport. Bacteria derived volatiles have also been found to suppress fungal growth. The bacteria *Bacillus subtilis* and *B. amyloliquefaciens* have been shown to reduce postharvest rot caused by *Penicillium spp. in vitro* by over 70% Arrebola et al. (16). Analysis of the volatiles produced by both *Bacillus* species found the most abundant compound was 3-hydroxy-2-butanone comprising over 50% of the volatile profile. In another study, the compound dimethyl disulfide was proposed as a potential biological control for the prevention of *Penicillium, Aspergillus*, and *Fusarium* postharvest rot after it was identified in the volatile profile of *Streptomyces* Wang et al. (15). Small volatiles produced by bacteria have shown promise as fungicide or fungistatic agents Schulz-Bohm et al. (17). For example, volatiles that repress growth from a distance could complement the physical barriers of rhizobacteria to prevent fungal root colonization Vespermann et al. (18), Neupane et al. (19), Kai et al. (20), Hoffman et al. (21). Unlike peptide effectors and larger metabolites (e.g., Tfe1,2, and prodigiosin), complex volatile profiles can function over larger distances (even several centimeters) Fiddaman and Rossall (22), Minerdi et al. (23), Garbeva et al. (24), Ossowicki et al. (25), Schmidt et al. (26). Although there is a clear role for bacterial metabolites in fungal biocontrol, few studies have identified the specific compounds involved in fungistatic interactions. (Table 1). In addition, environments, stage of life, and taxa influence volatile profiles Misztal et al. (27), which may influence long distance communication between fungi and bacteria Briard et al. (28), altering fungal behavior, morphology and biomass. Further characterization of bacteria and fungi chemical communication can help identify compounds that can have useful applications to medicine and agriculture.

**TABLE 1.** Studies documenting production of fungistatic volatiles by Serratia and Bacillus species.

| Inhibiting Organism | Inhibited Organism | Compounds Identified | Reference |
| --- | --- | --- | --- |
| *Serratia marcescens* | *Batrachochytrium dendrobatidis*,<br>*Batrachochytrium salamandrivorans* | No | Woodhams et al. (29) |
| *Bacillus megaterium* | *Aspergillus candidus*,<br>*Aspergillus fumigatus*,<br>*Penicillium fellutanum*, and<br>*Penicillium islandicum* | Yes | Mannaa et al. (30) |
| *Bacillus subtilis* | *Fusarium culmorum*,<br>*Neurospora crassa*,<br>*Trichoderma strictipile*,<br>*Verticillium dahliae* and<br>*Rhizoctonia solani* | No | Vespermann et al. (18) |
| *Bacillus subtilis* | *R. solani* | No | Fiddaman and Rossall (22) |
| *Serratia proteamaculans* | *Helminthosporium sativum*,<br>*Rhizoctonia solani*, and<br>*Sclerotinia sclerotiorum* | No | Popova et al. (31) |

Recent studies have identified a multitude of bacterial taxa with volatile-mediated fungistatic activity. However, bacteria-produced volatile profiles are complex, and progress in identifying individual components responsible for effects on fungi. To address this knowledge gap we explored volatile-mediated (indirect) antifungal effects by the bacteria *Serratia marcescens* and *S. proteamaculans*, and compared to *Bacillus subtilis*, known to produce inhibiting volatiles and used as a biocontrol species Fiddaman and Rossall (22). *Serratia* species are referred to in the literature as rhizobacteria Dhar Purkayastha et al. (32), meaning they can colonize the roots of host plants and confer advantages such as growth producing hormones and creating a physical barrier between a pathogen and the roots. *Serratia* is also known to produce volatiles containing compounds with effects on growth, antibiotic production and gene expression of neighboring fungi Garbeva et al. (24), Weise et al. (33), Schulz et al. (34). These understudied bacteria served as interesting models to study compared to the well-known, well-received biocontrol *Bacillus subtilis*. Most *Serratia*-produced volatiles have not been tested for inhibitory effects on fungal pathogens of plants, even though *Serratia* are commonly found associated with a variety of plant surfaces Ordentlich et al. (35), Dhar Purkayastha et al. (32). In this study, we show that fungal growth is deferentially inhibited by the volatiles of three different bacterial species, quantify differences in volatile emissions, and demonstrate the activity of specific volatiles against fungal pathogens. *Aspergillus fumigatus*, is an opportunistic pathogen and common genetic model organism. The Zoopagomycota fungus *Basidiobolus ranarum*, commonly lives in amphibian guts and can be an opportunistic human pathogen Khan et al. (36). *Mucor circinelloides* and *Actinomucor elegans* are zygomycetous fungi and opportunistic human pathogens. *Rhizopus stolonifer* is responsible for the postharvest rot of many fruits and vegetables. Finally, a genetic model organism and ascomycete, *Neurospora crassa* was evaluated. We employed six fungal species representing both diverse lifestyles and broad phylogenetic relationships. While our interest lies in the behaviour of many lesser known zygomycetous fungi, we attempt to validate our findings in popular model fungi (*N. crassa* and *A. fumigatus*. Our results suggest that some inhibition of fungal growth may be a result of indirect (volatile-mediated) mechanisms, contributing to our understanding of the biocontrol effects of *Serratia*.

## MATERIALS AND METHODS

### Strains

Volatile-mediated antifungal effects of three bacterial species were tested. *Serratia* species *Serratia marcescens* - Lab strain ADJS-2C_Red Aryal et al. (37) and *Serratia proteamaculans* - Lab strain BW106 Zhang et al. (38) were compared to a bacterial species with known indirect antifungal effects *Bacillus subtilis* - Strain E9 Na (39).

The bacteria volatiles’ fungistatic ability were tested on six diverse fungi listed in Table 2.

**TABLE 2.**
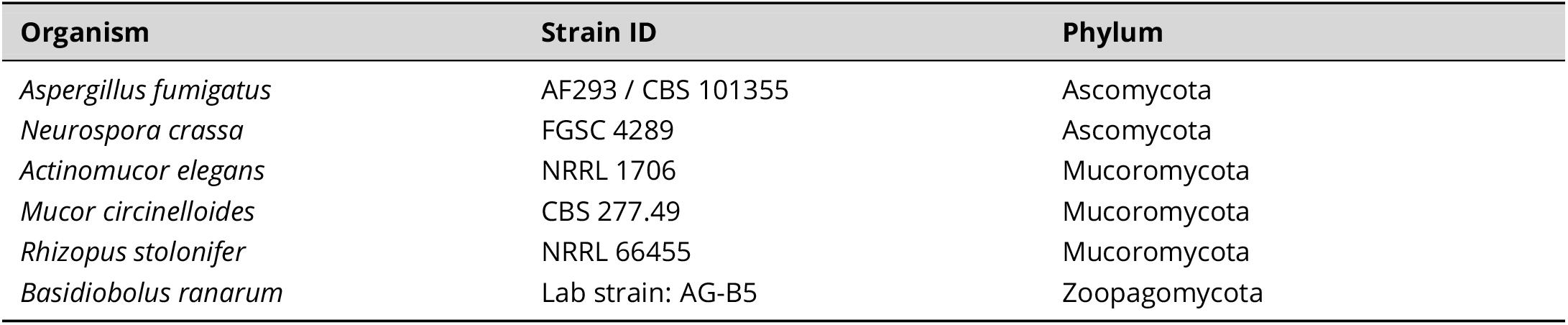
Fungal species evaluated for responses to bacteria-produced volatiles

| Organism | Strain ID | Phylum |
| --- | --- | --- |
| *Aspergillus fumigatus* | AF293 / CBS 101355 | Ascomycota |
| *Neurospora crassa* | FGSC 4289 | Ascomycota |
| *Actinomucor elegans* | NRRL 1706 | Mucoromycota |
| *Mucor circinelloides* | CBS 277.49 | Mucoromycota |
| *Rhizopus stolonifer* | NRRL 66455 | Mucoromycota |
| *Basidiobolus ranarum* | Lab strain: AG-B5 | Zoopagomycota |

### Inhibition assays with live bacterial cultures

Inhibition by volatiles were measured using a “donut plate assay” (hereafter “donut plates”) which consisted of a 100mm diameter Petri dish with a smaller 60mm Petri dish lid placed inside physically separating the bacterial media in the outside ring and fungal media on the inside ring. A volume of 15ml of Luria-Bertani (LB) media (10g Peptone, 10g NaCl, 5g Yeast Extract, 7.5g Agar, and 500ml Water) was pipetted into the outer ring of a 100mm Petri dish with a 60mm Petri dish lid placed inside. Then 8ml of Malt Yeast Extract Agar (MEYE) (1.5g Yeast Extract, 1.5g Malt Extract, 5g Dextrose, 7.5g Agar, 500ml Water) was added to the 60mm Petri dish lid. *N. crassa* was grown on Vogel’s media http://www.fgsc.net/methods/vogels.html. To inoculate the outer ring with bacteria, a single colony picked with sterile toothpick was inoculated into 25 ml liquid LB in 50ml conical tube, shaken overnight at 100 RPM at 28°C). Growth rates of the bacteria were compared by measuring optical density every 30 minutes using Corning®96-well Clear Flat Bottom Polystyrene (Product Number 3370) with 200 *μ*l LB media and 1 *μ*l initial inoculum cell concentration for *Serratia* or *Bacillus* was measured at the absorbance at 595 nm and 700 nm to avoid interference of the red prodigiosin compound produced *S. marcescens* Haddix et al. (40). Absorbance for cultures of *B. subtilis*, and *S. marcescens* grown overnight (n=9) were comparable (Supplemental Data - Table 1). One milliliter of media from these cultures was innoculated onto the outer LB ring and allowed them to grow for either 24 or 48 hours at 28°C before introducing the fungi. At 48 hours, fungal growth was equally inhibited by all *Serratia* and *Bacillus* strains tested, so this was used as the default time for bacteria to grow before fungal inoculation.

Fungi were grown to sporulation/conidiation (1 week) at which point the spores were collected with a sterile toothpick then stored in a 1.5ml microcentrifuge tube with 1ml of autoclaved water at 4^°^C. *Basidiobolus* do not sporulate under lab conditions, therefore 1cm × 1cm mycelial plugs were harvested from one-week-old MEYE plates and stored in autoclaved water at 4°C. Mycelial plugs (*Basidiobolus*) or 1000 spores/conidia (*Actinomucor, Mucor, Rhizopus, Aspergillus, Neurospora*) were added to the media in the central 60mm Petri dish lid and incubated at 25°C. The diameter of fungal mycelia were measured 24, 48, 72, and 96 hours after fungi were inoculated. Each experiment had three technical replicates of one to three biological replicates for each condition.

### *In vivo* growth inhibition

Strawberries (Monterey cultivar) were grown at UCANR South Coast Research and Extension Center, with no fungicides. Strawberries were cut and divided to quantify starting colonization levels (one half) and evaluate fungistatic effects of volatiles (second half). The fruit was inoculated on top of the skin with approximately 1000 spores of *R. stolonifer* and incubated at 28°C for 48 hours in 12-hour day/night cycles. To quantify initial fungal growth, each strawberry half was placed in a 50 mL conical tube with 25 mL of water and homogenized. The homogenized mixture was allowed to settle for 5 minutes, long enough for the strawberry pieces to fall out of solution, while spores remained suspended. The supernatant was collected and spore count was quantified with a hemacytometer method (described above) by taking the average of three technical replicates of each sample. The second half of each strawberry was placed inside a donut plate with 48-hour old culture of *S. marcescens* in the outer ring. Strawberries remained on the plates for 48 hours followed by fungal growth quantification using the spore count by hemacytometer. Each growth and control condition was tested in at least three replicates.

### Quantification and identification of bacterial volatiles

A liquid LB media flask was inoculated with a bacterial strain and grown overnight as described above. From the overnight culture, 1 ml (OD 700 - 1.5) was used to inoculate plastic 60 mm Petri dishes and incubated for 48 hours at 28°C. Volatiles from these 48 hour bacteria cultures were collected using a pull-only collection system. *B. subtilis* was used as a positive control due based on previous work which demonstrated the bacteria can produce volatiles with antifungal activity. Volatiles were also collected from empty Petri dishes and Petri dishes with LB media as negative controls. Petri dishes were enclosed in 350 ml Mason Ball jars with airtight Teflon lids having two Swagelok connection ports. One port was fitted with a charcoal filter (copper pipe filled with activated carbon) to remove odors from incoming air, while the other was fitted with an adsorbent trap (0.25 inch glass tube filled with 40 mg HayeSepQ beads 80-100 mesh size) to collect volatile emissions. On the end of the adsorbent trap a vacuum hose was attached and air pulled through the trap at a rate of 0.5L/min for 6 hours at 25°C. Volatiles were eluted into vials by passing 150 *μ*L dichloromethane spiked with 4ng/μL nonyl-acetate, and 2 ng/*μ*L octane through each trap and pushing into the vial using a gentle stream of nitrogen. Samples were stored at −80°C until analysis by gas chromatography and mass-spectrometry. In total, nine collections were performed for each species (n=9). Samples were analyzed using a Thermo FisherTRACE 1300 gas chromatograph (GC) linked to a TSQ Duo Triple Quadrupole mass spectrometer (MS) operating in a single quadrupole mode. The GC was fitted with a 30m TG-5MS column (Thermo Fisher), 0.25 mm diameter with a stationary phase of 0.25 *μ*m. The inlet temperature was set to 220 °C and operated in splitless mode. Helium was used as a carrier gas delivered via constant flow at a rate of 1.2 ml/min. The transfer line to the MS was held at 280 *°*C and the ion source was operated at 250 °C. The instrument was tuned to proper settings for electron ionization mode using the autotune feature. MS detection was performed by scanning atomic masses from 30-500 at a scan rate of 0.2 seconds. One microliter of the sample was injected using an autosampler, volatilized in the inlet, and recollected on the column, which was held at 40 °C for one minute following injection. Following this, the column temperature was increased linearly by 8 C/min up to 280 °C, then held for one minute, after which data recording for that sample was terminated. The instrument was cooled to 40 °C for the next sample run. Spectral outputs were evaluated using Chromeleon 7 software. For all experimental samples (*Serratia* species and *B. subtilis*) Microsoft Excel table outputs of each peak retention time and area were generated along with putative identifications based on comparison to spectra in the National Institute of Standards and Technology (NIST) library. These results were compared to outputs for both negative controls to identify compounds originating from the Petri dish and media, both of which could contribute volatiles. Matches between negative control contaminants and peaks in the experimental samples were confirmed by spectral comparison. Any trace contaminants from column and septum bleed were also removed. The reduced list of compounds emitted by experimental treatments was then examined for matches to known spectra in the NIST library. Compounds with Reverse Match Factors (RSI) of 85% or higher, and which were released consistently across samples for a given species, were retained for further quantification, analysis, and examination as pure compounds in various concentrations.

### Volatile quantification and analysis

Peaks in the total ion chromatogram were integrated and areas were used to calculate the total quantity of volatile present in the entire sample (representing compound sampled over the 6 hour collection period). Calculations were performed relative to the internal standard (nonyl acetate) peak area and concentration (4ng/*μ*L). Compounds with less than RSI of 850 (85% confidence prediction) were not included in further analyses due to logistical constraints on pursuing their identification. The list of compounds detected and their amounts can be found in Supplemental Table 2. Compound IDs and amounts from this table were used for subsequent multivariate analysis with the R package, MetaboAnalyst 4.0 (R version 3.6.0). MetabolAnalyst analyses compared volatile emissions among treatments of the entire volatile profile, as well as examining individual compounds. Auto-scaling centered around the volatile production average was used to make all metabolites comparable. Analysis of Variance followed by posthoc Tukey tests identified volatiles emitted in significantly different quantities among the bacterial strains. Non-metric multidimensional scaling (NMDS was performed with the function metaMDS from the vegan package. Code used for analysis with MetaboAnalyst and NMDS are archived in the Github repository (https://github.com/stajichlab/Serratia_Bacillus_Volatiles_Analysis).

### Volatile effects on Vacuolization of Fungal Hyphae

To test the direct impact of volatiles on fungi, *R.stolonifer* colonies were growth by inoculating 1000 spores suspended in autoclaved water onto 60 mm Petri dish with 8 ml MEYE media and agar. The spores were allowed to germinate and colonies grow for 12 hours at 25°C. Control colonies were allowed to grow for an additional hours. Treatment colonies were exposed to volatiles for 1 hour. Volatile were applied by placing 10*μ*l of compound on a 1cm × 1cm piece of filter paper approximately 2 cm away from the *R. stolonifer* mycelium. Hyphae were excised from the media with a sterilized metal scalpel and imaged on microscope slide on an Amscope Compound Microscope with a 40X Phase Objective lens.

### Growth inhibition by volatiles

The compounds Tropone(Sigma-Aldrich 252832), 5-Methyl-2-furyl methanol (Sigma-Aldrich CDS003383), Lepidine (Sigma-Aldrich 158283), 2,5-dimethylpyrazine (Acros Organics AC174520050), 2-undecanone (Sigma-Aldrich U1303), Anisole (Acros Organics AC153920050), and dimethyl trisulfide (Tokyo Chemical Industry Company D3418), were tested for growth inhibition on fungi. 10mg of the compound were pipetted onto a 1cm × 1cm filter paper in the outside ring of a donut plate with 1000 spores of *R. stolonifer* on MEYE in the central Petri dish. After 24 hours the fungal mycelium diameter was measured. Each compound concentration was tested with three independent growth plate replicates.

## RESULTS

Fungal colonies grown with *S. marcescens* demonstrated reduced growth rates compared to the unexposed control (streak plate Figure 1A). The difference in colony size was quantitatively assessed by measuring colony width and shape for cultures with and without bacteria streaked on either side. Streak plates demonstrate bacteria can inhibit fungal growth without direct contact, however the mechanism could be indirect via diffusion of bacterial metabolites or proteins in the growth medium or through air-borne volatilized compounds. To evaluate these possible routes, we developed the donut plate assay where fungi and bacteria were grown on two separate Petri plates which were allowed to share common headspace but not a common medium. This set-up allows for the exchange of gases between the fungi and bacteria, prevents direct contact, and allows each microbe to be grown its preferred or specialized growth medium (Figure 1B-D).

**FIG 1.**
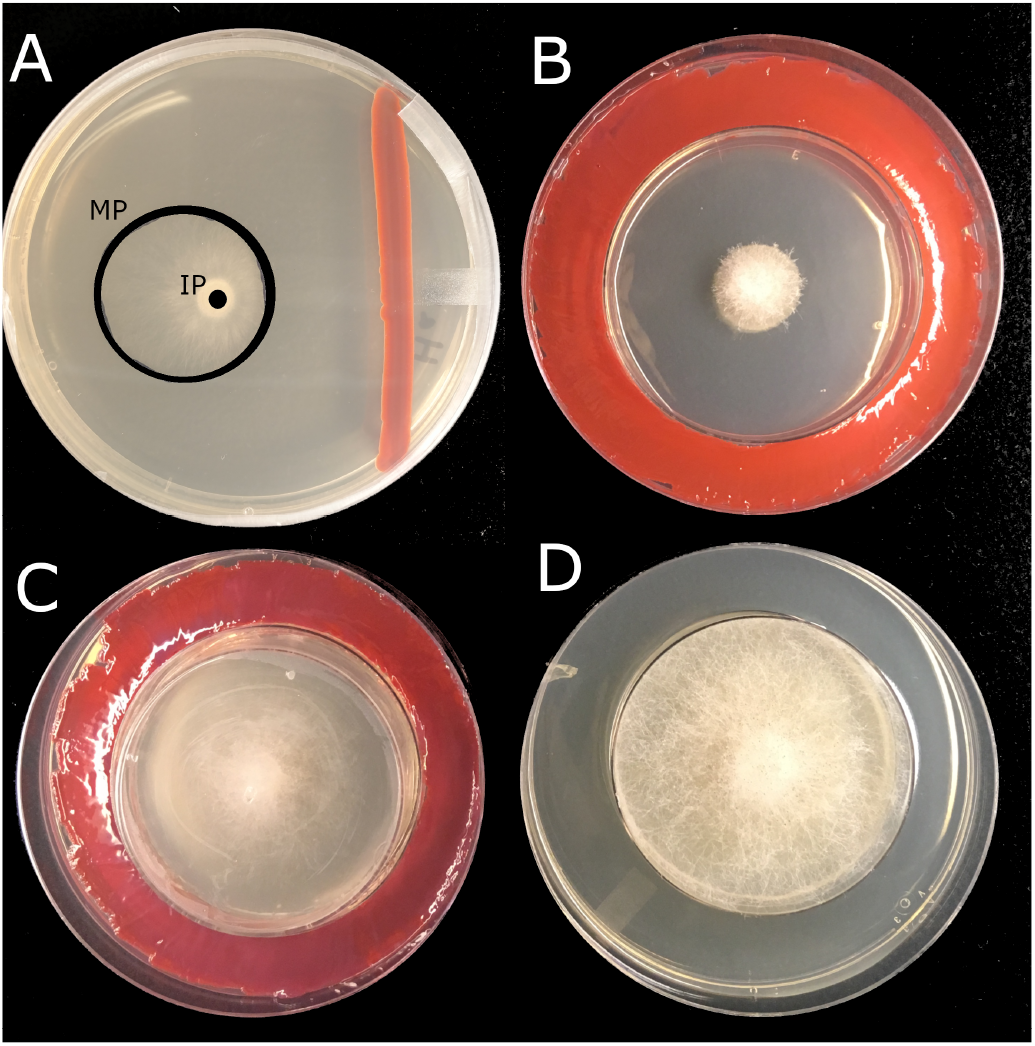
Donut plate assay reveals gases are responsible for fungal growth repression. A. *R. stolonifer* grown for 48 hours with *S. marcescens* streaked on one side. IP=Innoculation Point, MP=Mycelium Perimeter. B. Donut plates with *R. stolonifer* inside. C. *R. stolonifer* covered preventing volatiles from entering the inner plate. D. Normal growth of *R. stolonifer* without *S. marcescens*. For each condition n=3

Testing if fungal growth was impacted equally by the presence of each bacterial isolate (Table 2) we found that *Serratia proteamaculans* imparted the strongest growth inhibition followed by *S. marcescens* and *B. subtilis*.The bacteria colony density was kept constant by first confirming that *Serratia* and *Bacillus* strains required 24 hours of growth to reach stationary growth phase (supplemental data table 1, supplemental figure 1). Subsequent donut plate assays demonstrated that volatile exposure was sufflcient to inhibit growth.

In the first 24 hours of exposure to volatiles from *S. proteamaculans, N. crassa* colonies were about 10% the size of the control with no volatile exposure. *R. stolonifer* colonies were about 15% of its control size at 24 hours when incubated with *S. proteamaculans*. However we noted that *A. elegans* grew under these conditions with no detectable inhibition. *M. circinelloides* had an intermediate level of inhibition and achieved 30% of the colony growth seen in the unexposed control. *N. crassa* growth was also repressed by the other two bacterial volatile profiles; colonies were significantly smaller at the 24 hour time point (Figure 2). *R. stolonifer* was also inhibited by volatiles of all strains at 24 hours, but only *S. proteamaculans* and *S. marcescens* significantly inhibited growth at 48 hours (Figure 2). *M. circinelloides* was inhibited by volatiles of *B. subtilis* at 24 hours (Figure 2). The growth of the Zoopagomycota fungus *B. ranarum* was inhibited by volatiles of all bacterial species at 48 hours (Figure 2). The Mucoromycotina fungus *A. elegans* was resistant to the volatiles of all tested bacteria (Figure 2), showing little growth repression. The growth of ascomycete *A. fumigatus* was inhibited at 72 hours (Figure 2) by all three bacteria species’ volatiles. Overall *A. fumigatus* has the slowest growing hypha as the mycelium coveres only about 400 mm^2^ (20% of the plate) after 72 hours.

**FIG 2.**
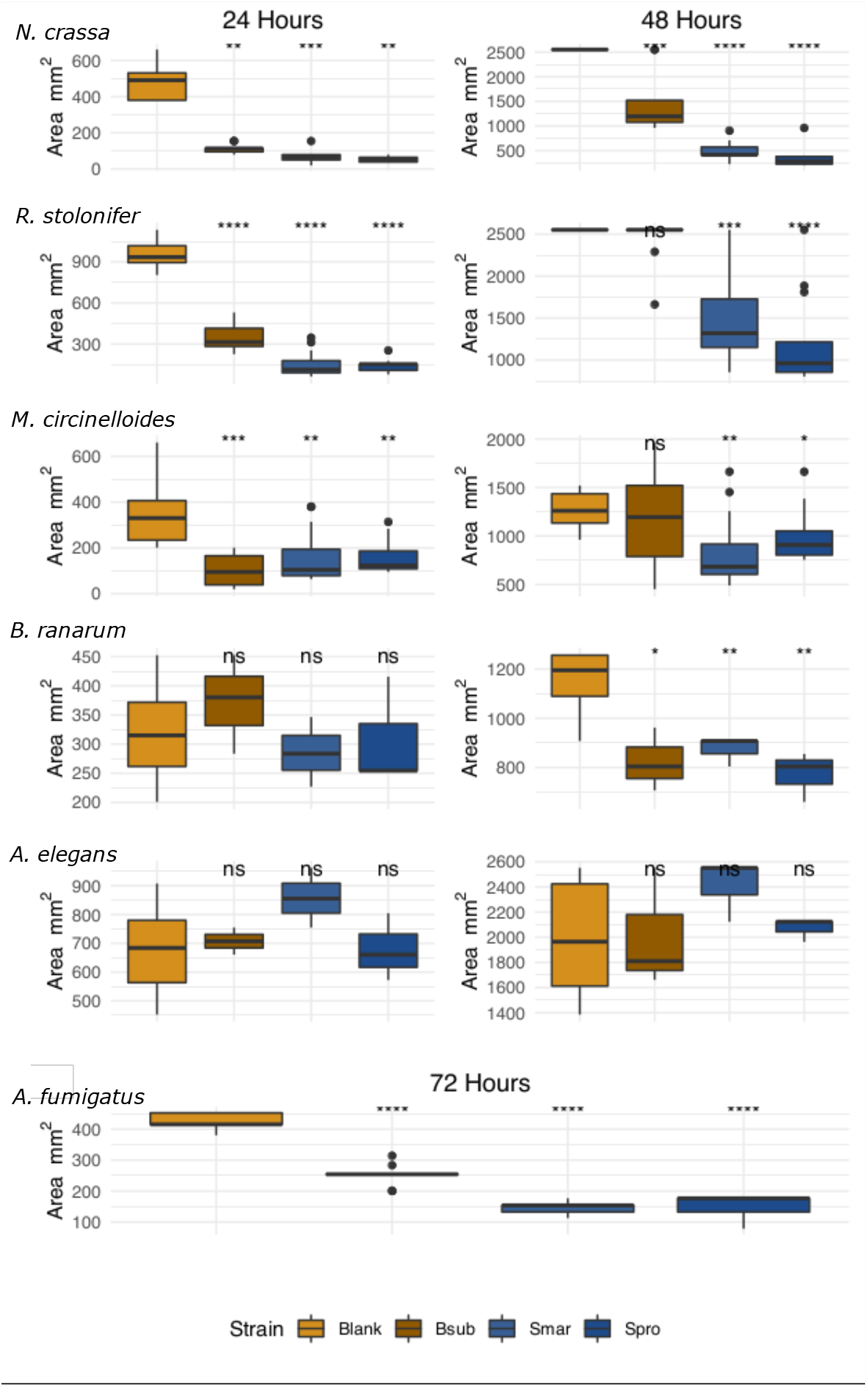
Five of six tested fungi show significant growth reduction compared to fungi in a donut plate without bacteria. Growth of *N. crassa, R. stolonifer, M. circinelloides, B. ranarum, A. elegans* after 24 hours (right) or 48 hours (left) with and without bacteria volatiles, or *A. fumigatus* after 72 hours. n>=3

To determine the effectiveness of the compounds to prevent postharvest rot, we incubated strawberries inoculated with *R. stolonifer* in the presence of *S. marcescens*. After incubation on strawberry and average of 100 spores/*μ*l of *R. stolonifer* were recovered, however, on the strawberries exposed to *S. marcescens* volatiles during the infection period only an average of 36 spores/*μ*l were recovered. (Figure 3).

**FIG 3.**
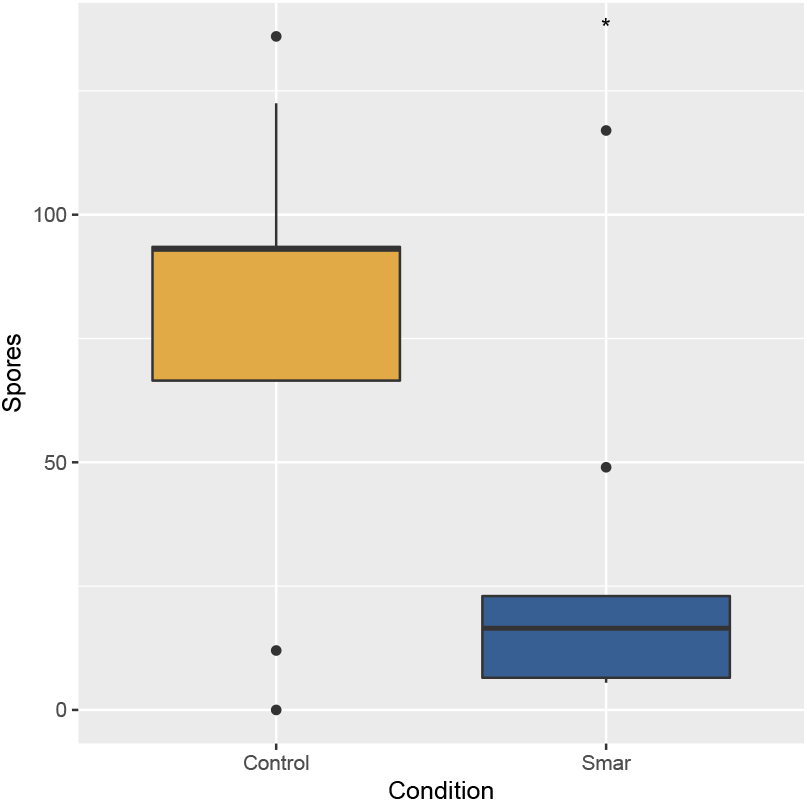
Strawberries grown with *S. marcescens* (Smar) had significantly fewer spores of *R. stolonifer* recovered compared to the untreated condition. * - p-value < .05

Analysis of the volatilized compounds produced by the bacteria identified 62 compounds produced in common to all strains. An average of 500 ng of total volatiles were collected from each culture during the sampling period. Samples which recovered less than 200 ng of detected metabolites were removed from the analysis (Figure 4A) and were (*B. subtilis* A, G and I, *S. marcescens* D, F, G and *S. proteamaculans* I. The volatile profiles were distinctly different for the two examined bacteria genera. *B. subtilis* produces a unique set of compounds that are absent in the *S. marcescens* and *S. proteamaculans* volatiles, however the two *Serratia* isolates had qualitatively similar profiles. Quantitative comparison of the *S. marcescens* and *S. proteamaculans* profiles does find differences in the amount of each compound produced (Figure 4B-D). For example, dimethyl trisulfide is the most abundant compound in the *S. proteamaculans* profile, while anisole was abundant and unique to *S. marcescens. B. subtilis* produces multiple volatile compounds at high abundance, including 2-undeconol/2-undecanone, and butanoic acid.

**FIG 4.**
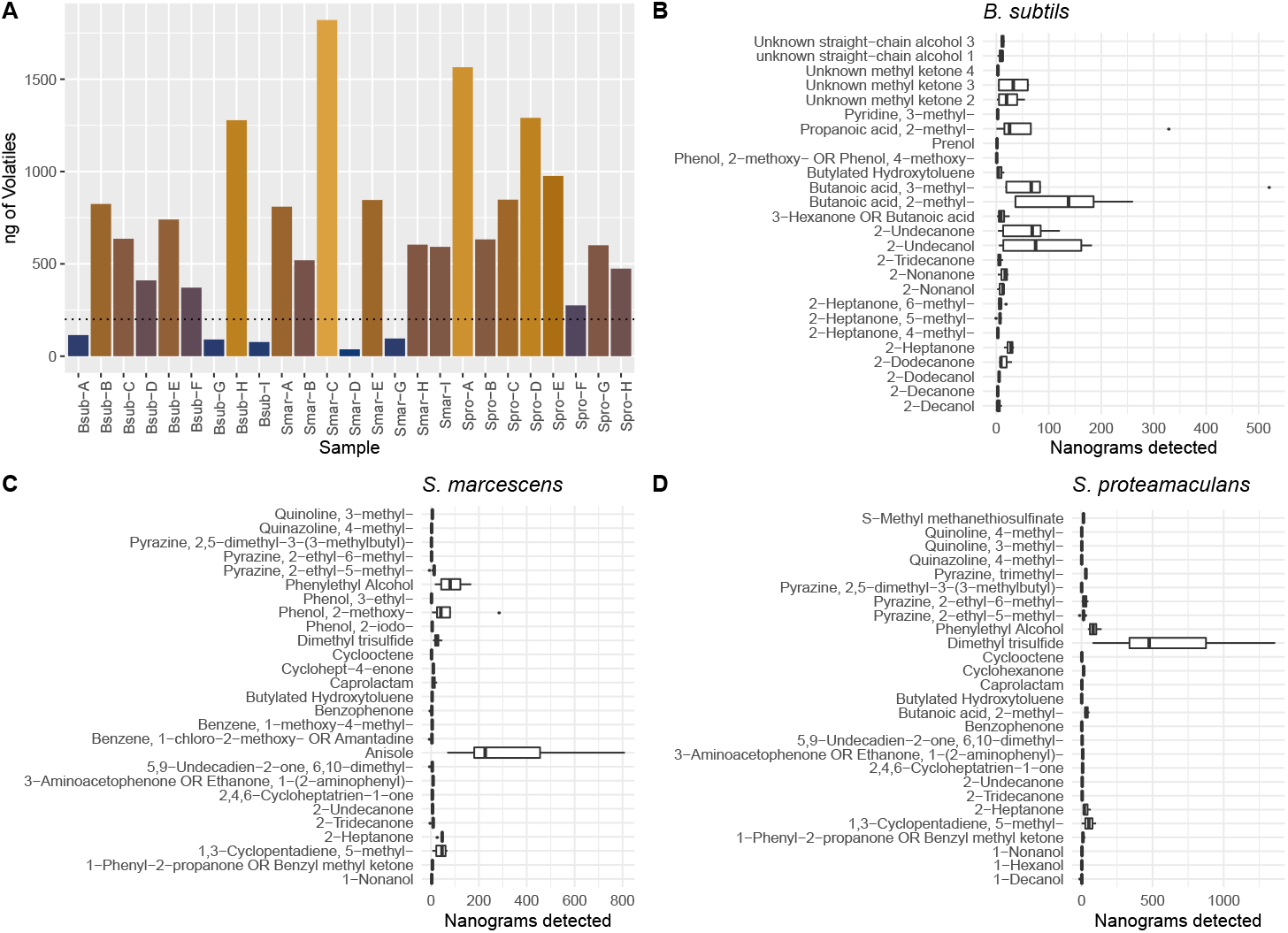
GCMS output: Total measured product detected from samples in nanograms. 25 out of 27 samples had greater than zero nanograms of metabolites detected. A. Total output per sample. Horizontal line at 200 ng represents cutoff requirement for sample to be included in volatile profile comparisons. Bsub = *B. subtilis*, Smar = *S. marcescens*, Spro = *S. proteamaculans*. B. The sum of volatiles produced by all samples of one species *B. subtilis* C. *S. marcescens* D. *S. proteamaculans*

We classified compounds detected in 4 or more replicates for a species as a robust and reliable component of the volatile profile resulting in a total of 62 compounds found in three species. *B. subtilis* produces 18 unique compounds, which are not detected in the other two bacteria. There were 4 compounds in *B. subtilis* profile that were also found in both of the *Serratia* species.The *S. proteamaculans* volatile profile had 4 unique compounds and 11 also found in *S. marcescens. S. marcescens* had 7 unique compounds (Figure 5A). Analysis of the compound profiles using Non-Dimensional Multivariate Scaling (NMDS) shows clear separation of each of the bacteria volatile profiles which is due to differences in composition and amount of volatile produced. (Figure 5B)

**FIG 5.**
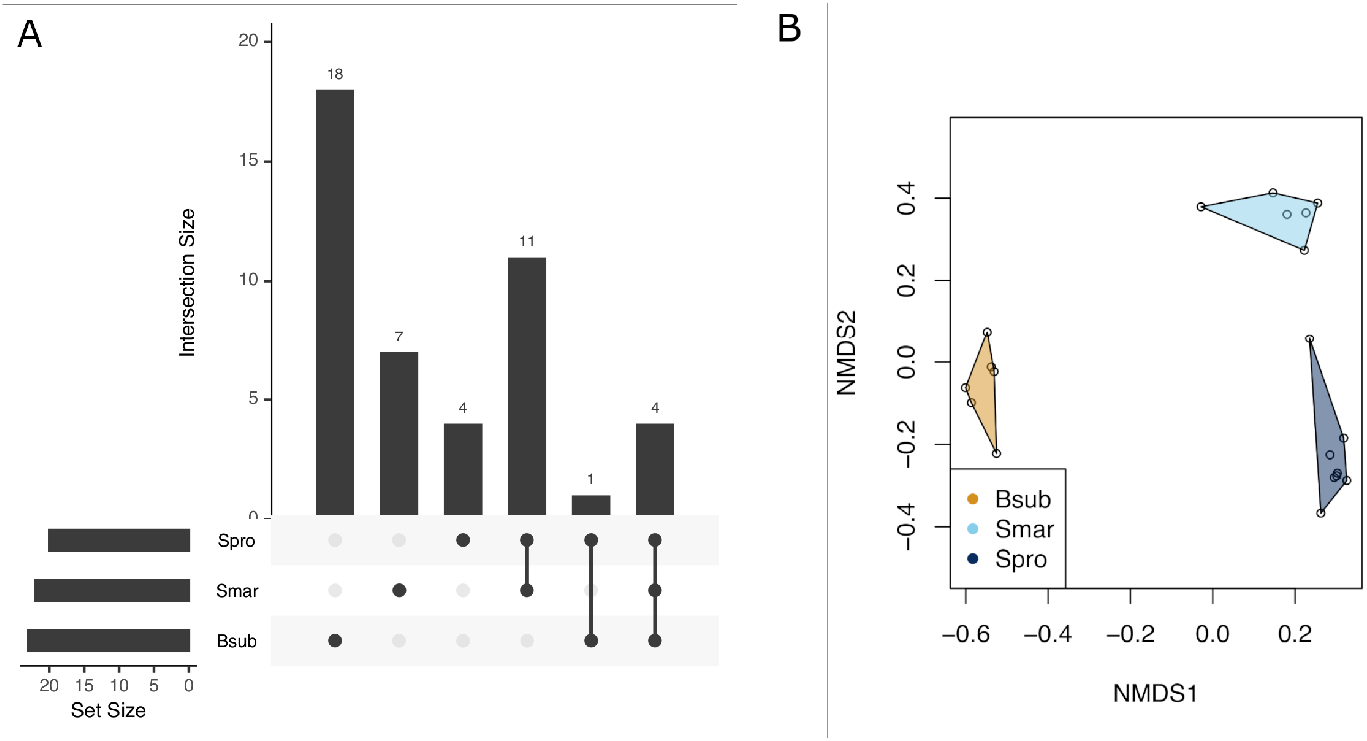
Comparison of the volatiles of *Serratia* and *Bacillus*. A. Upset plot showing overlap of common volatiles detected between the three species. B. Bacterial volatiles amounts variation and compound presence/absence shows three distinct profiles in non-dimensional scaling.

The relative and approximate abundance of volatile compounds was predicted by incorporating an internal standard. Comparisons of the *B. subtilis*, *S. marcescens* and *S. proteamaculans* measurements were performed to test what compounds could differentiate the species. An ANOVA analysis (*p*-value < .05) and Tukey’s/HSD analysis revealed 22 volatiles compounds produced at significantly different abundances 6. Dimethyl trisulfide had of the highest p-values from ANOVA analysis. Upon further inspection of the normalized concentrations of dimethyl trisulfide in all the bacteria samples, we show that *S. proteamaculans* was the only significant producer. Anisole was another compound that only had one bacteria producing it, *S. marcescens*. 2-Undecanone was produced at very low levels in the *Serratia* species but was produced at much higher levels in *B. subtilis*. Our ANOVA results indicated 2-undecanone was present at significantly increased levels in *B. subtilis* profile. Other samples had significant differences but these were the result of measured reductions of a specific compound compared to the other two bacteria. Specific p-values and comparisons are found in Supplemental Table 2.

The ANOVA identified several compounds in the bacteria volatile profiles that we hypothesized could be responsible for fungal growth inhibition such as 2-undecanone, dimethyl trisulfide, anisole. Furthermore, 2-undecanone was chosen based on previous work demonstrating its activity as an antifungal Popova et al. (31). It is also used in food, fragrance, and is an effective insect repellent approved for human use Bohbot and Dickens (41). Dimethyl trisulfide was selected for further testing based on the abundance in the *S. proteamaculans* profile. Finally, anisole was selected as it was uniquely and abundantly produced by *S. marcescens*. We also selected tropone, lepidine, 2,5-dimethylpyrazine for further testing based on their high abundances. All of these compounds were tested for growth inhibition activity on the fungus *R. stolonifer*.We found that 10 mg of volatilized 2-undecanone, anisole, and dimethyl trisulfide were able to inhibit growth (Figure 7).

**FIG 6.**
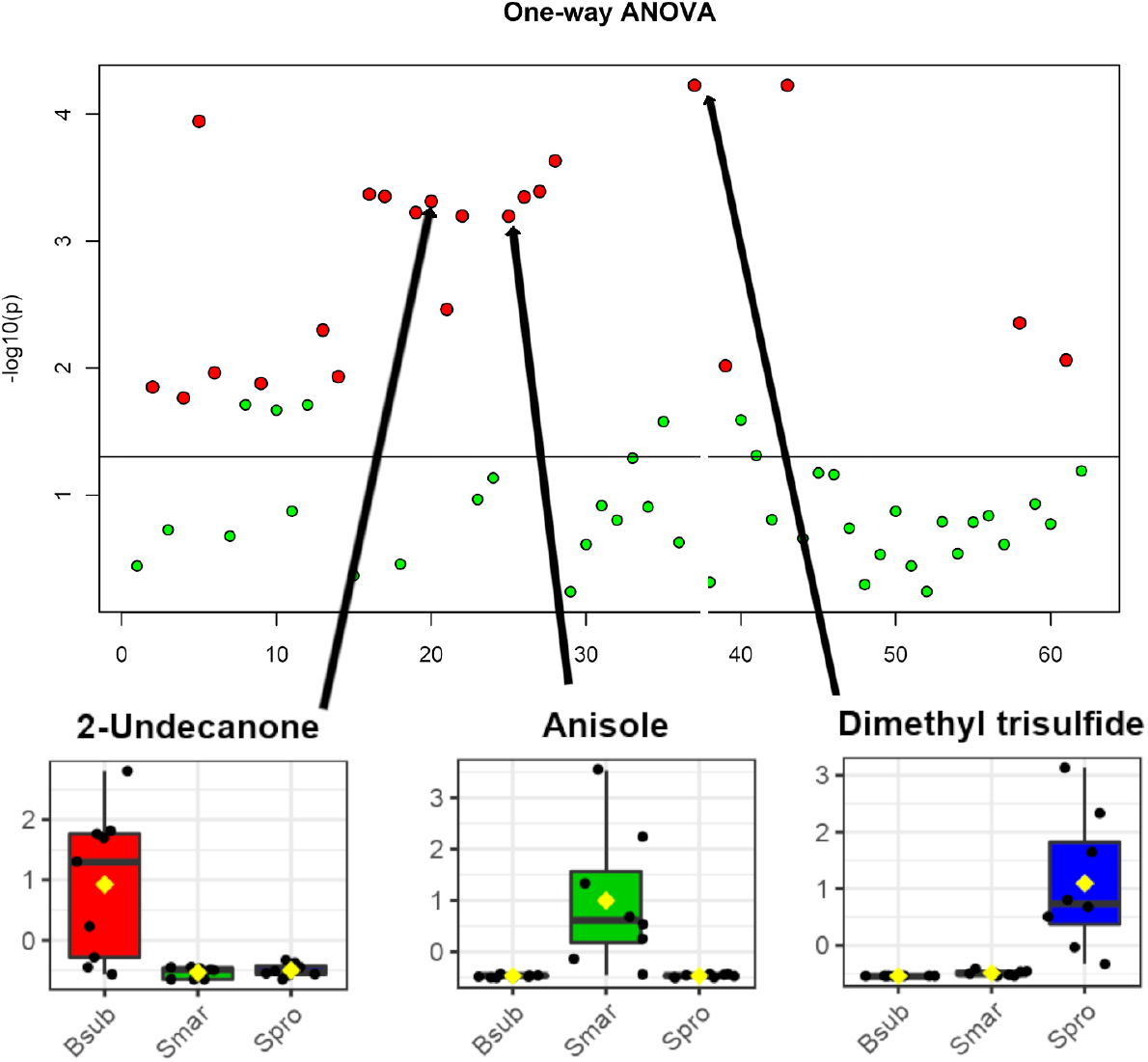
ANOVA analysis of differential volatile production by each bacterial species. Each red dot represents comparisons of one volatile among the three bacteria with significant differences between species. 2-Undecanone is production is highest in *B. subtilis*,anisole in *S. marcescens*, and dimethyl trisulfide in *S. proteamaculans*.

**FIG 7.**
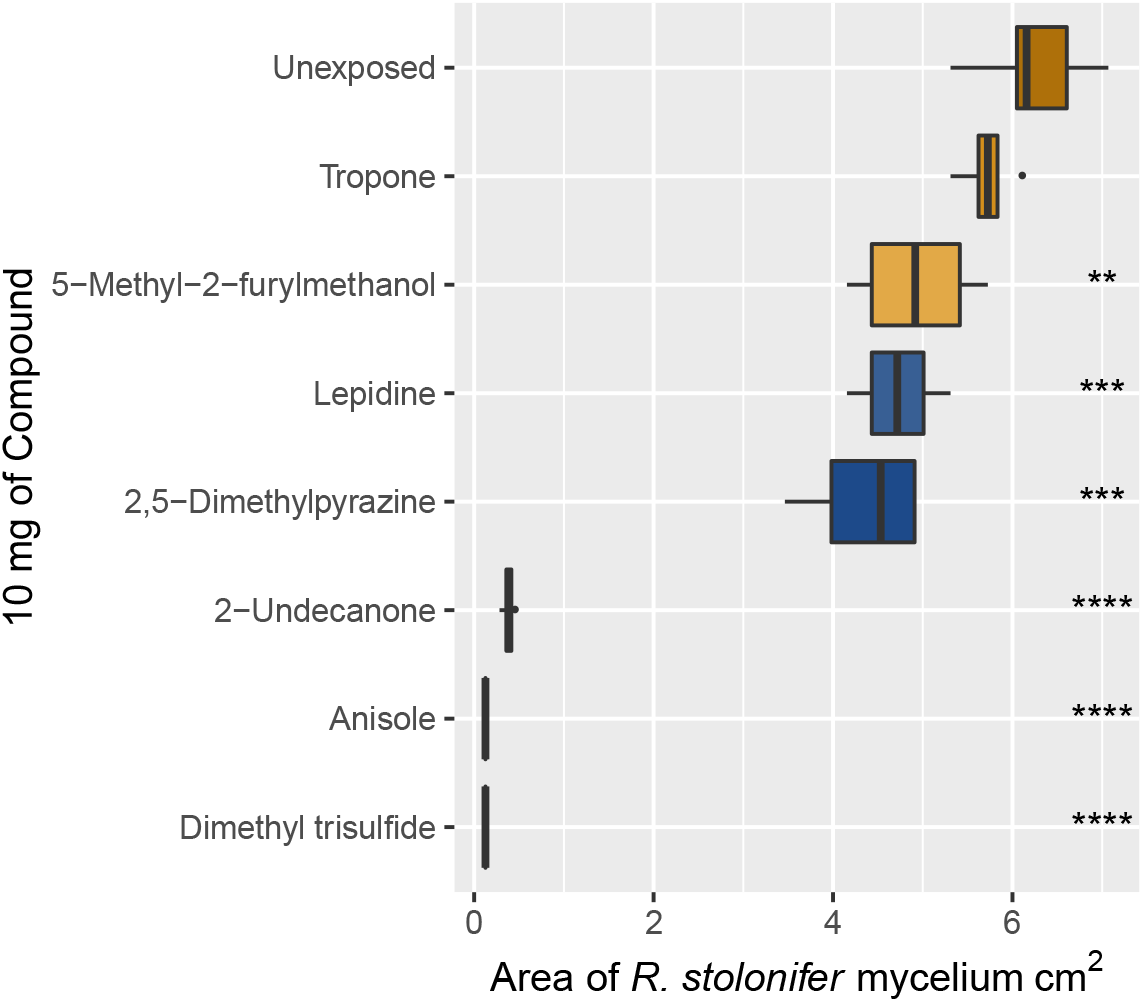
Growth inhibition of *R. stolonifer* with a single volatile compound. Anisole, dimethyl trisulfide, and 2-undecanone inhibit fungal growth as volatiles compounds. A. *R. stolonifer* germination and growth from spores in the presence of 10 mg of select bacterial volatiles. n=3 B. Strawberries grown with *S. marcescens* had significantly fewer spores of *R. stolonifer* compared to the untreated condition, * - p-value < .05, **=.01, ***=.001, ****=.0001

Increased vacuole visibility and cytoplasmic irregularity was observed in fungal hyphae exposed to anisole and dimethyl trisulfide (Figure 8), but not those exposed to 2,5-dimethylpyrazine, 2-undecanone, or the control with no volatile exposure. We did not observe vacuoles or granulated cytoplasm in microscopic observations of the control hyphae which were not exposed to volatiles. The vacuole becomes very pronounced and the cytoplasm lost transparency and formed dark, granulated structures in the hyphae exposed to anisole and dimethyl trisulfide (Figure 8D and E). This vacuolization and cytoplasmic thickening has been noted in previous work involving the fungal response to bacterial volatiles. In *Sclerotinia sclerotiorum* hyphae exposed to volatiles of rhizobacteria induced vacuole formation, condensed cytoplasm possible due to loss of cell membrane integrity Giorgio et al. (42). In the anisole and dimethyl trisulfide treatments, approximately 45-70% of the hyphae showed pronounced vacuoles or condensed cytoplasm after only one hour (supplemental figure 1 and supplemental table 4).

**FIG 8.**
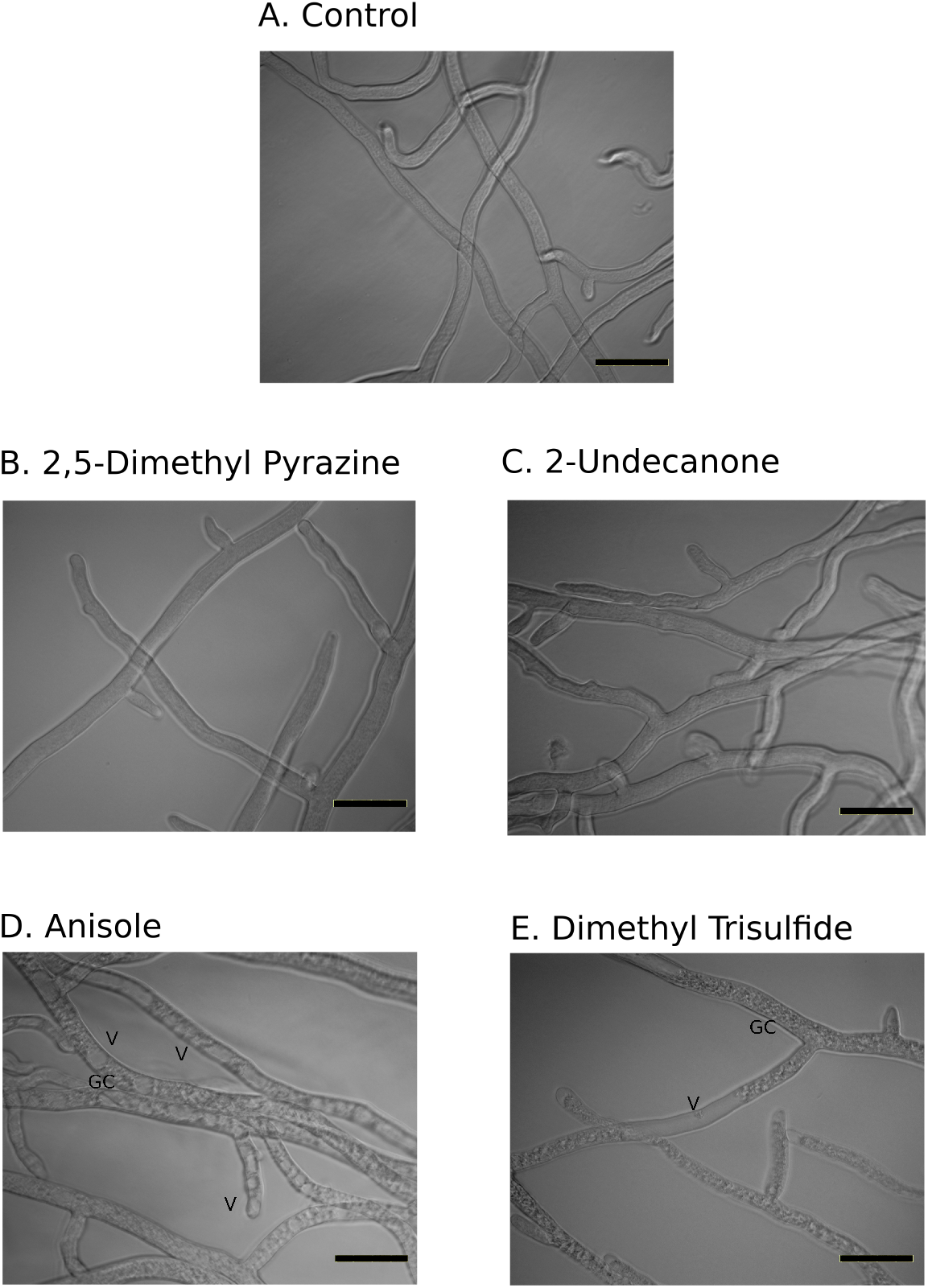
Microscope images of 1 hour of volatile exposure to hyphae of *R. stolonifer*. A. Unexposed to volatiles, normal growth. B. Dimethyl-pyrazine exposed hyphae. C. 2-Undecanone D. Anisole exposure results in moderate vacuolization E. Dimethyl trisulfide vacuolization. V - Vacuole, GC = Granulated cytoplasm. Bar=.2 mm. Each condition had 3 biological replicates, images are representative of findings

## DISCUSSION

Our observation of the growth inhibition potential of bacterial volatiles on fungi led to the identification of bioactive compounds, analysis of differences between bacterial species, and demonstrated fungistatic potential of pure compounds. The novel donut assay, specifically developed to study the effect of headspace, demonstrated that diverse fungi were susceptible and growth inhibition was observed for most strains. Additionally, we show that *Serratia* species are capable of producing two compounds in relatively high amounts that are fungistatic to *R. stolonifer* a postharvest rot of many fruits and vegetables. These results indicate *Serratia proteamaculans* and *Serratia marcescens* should be further investigated for their biocontrol potential.

The rhizobacteria used here are only a few of many that naturally occur in soil. The wide range of volatile products found here and the diverse range of fungal species they affect illustrate the vastness of products that can come from studying bacteria fungal interactions. *R. stolonifer* showed dramatically reduced colony growth in the first 48 hours when exposed to volatiles from each of the three bacterial species 2. *M. circinelloides* was less inhibited than *R. stolonifer* but still had significant growth reduction (average control area 300 mm^2^ versus 100 mm^2^ of volatile exposed samples). The slower growing *B. ranarum* showed significant inhibition only in the second time point (48 hours) but not in the first 24 hours. This delay may also be a result of the inoculation method used, as *B. ranarum* was grown from hyphal plugs instead of using spores like the other fungi tested. This may indicate that spores may confer some level of protection by delaying germination in the presence of bacterial volatiles, but more testing is needed to verify this. Interestingly, the Mucoromycotina fungus *A. elegans* never showed significant growth inhibition when exposed to volatiles. This observed resilience warrants further investigation to understand molecular or physical properties of this fungus that prevents it from being impacted by the compounds. *A. fumigatus* was the slowest growing organism tested and showed measurable growth inhibition upon exposure to volatiles of each of the three bacteria at the 72 hour time point. All of these observations were made using the donut plate assays where the fungi, growing on rich media, were exposed to the bacteria volatiles for 48 to 72 hours. We further explored the inhibition of *R. stolonifer* on a more realistic growing surface, strawberries. The strawberry is a natural substrate for *R. stolonifer* which may affect the germination of spores, growth of the hyphae, and resistance to bacteria volatiles. We showed that 48 hours of *S. marcescens* volatiles have the capacity to inhibit growth of *R. stolonifer* on fresh strawberries, demonstrating its effectiveness in semi-realistic environments.

Many publications have highlighted the phenomenon of bacterial volatile effects on plants and fungi, but few go on to identity the compounds. To address this, we attempted to identify many of the compounds in the volatile profile using gas chromatography and mass spectrometry. We characterized volatile blend compositions and identified species-specific compounds that have potential use as tools to inhibit fungal growth. For example, anisole was found in seven out of eight samples of *S. marcescens* at amounts greater than 50 ng and was not detected in the volatiles of the other bacteria (Supplemental Table 3). 2-Undecanone was found in 7 out of 9 samples of *B. subtilis* at approximately 10 fold higher amounts than in *S. marcescens* or *S. proteamaculans*. Dimethyl trisulfide was produced at a remarkably high level, on average over 600 ng/sample of *S. proteamaculans*. These specific differences all contribute to the spacial differences we noted in the NDMS plot. The three bacteria profiles clearly differentiate and show no overlap indicating that all three of our tested profiles are different. These results reveal that even closely related bacterial species can produce very different volatile profiles, both qualitatively and quantitatively, with significant implications for biocontrol potential.

Even with multiple fungistatic compounds found in the different bacterial volatile profiles, our experiments document important ranges in the sensitivities of different fungi to volatile-based inhibition. Across all experiments, *N. crassa* and *Aspergillus* (both ascomycetes) were more susceptible to the volatiles produced by the bacteria than the Mucoromycotina fungi *R. stolonifer* and *M. circinelloides. A. elegans*, also a member of the Mucoromycotina, demonstrated a robust resistance to the volatiles suggesting further study is needed to understand how these fungi process or avoid the compounds. Additional research is needed to identify if taxonomic or genomic content correlates to fungal sensitivity or resistance to inhibitory volatiles from bacteria. The experiments presented here with a limited number of representative species raise the possibility that certain groups (e.g., Ascomycota) may be more susceptible to fungistatic volatiles than others. The variation in sensitivities observed in the fungi may be a result of relative frequency that these groups are competing with bacteria. If factors can be identified that modulate fungal resistance it will be informative to evaluate the evolutionary history of these components to better understand their dynamics. Likewise, a more thorough profile of bacterial species to understand the changes in volatile production in response to biotic and abiotic stimuli would help determine whether these pathways are actively regulated as part of microbial interactions.

Among the volatile compounds emitted by cultures of the bacterial species studied here, several were produced both consistently and at a high abundance. These include 2-undecanone and other long-chain fatty acids, anisole, dimethyl trisulfide, and butanoic acid. A surprisingly high volume of several compounds were recovered over a six hour collection time. 2-Undecanone had an average of over 60 ng per sample. Anisole recovery was over 200 ng per sample. However, dimethyl trisulfide had an average of almost 500 ng recovered per sample. Of the compounds tested for fungistatic properties, 2-undecanone, anisole, and dimethyl trisulfide had the strongest activity. The compounds anisole and dimethyl trisulfide appear to induce vacuolization of hyphae indicating a possible induction of stress on the fungal cells. This phenotype was observed in the ascomycete *Colletotrichum* when exposed to dimethyl trisulfide (43) and *Sclerotinia sclerotiorum* exposed to dimethyl trisulfide (at minimal inhibitory concentration of 24 milligrams as the headspace source) and 2-undecanone (15 milligrams as the headspace source) Giorgio et al. (42). Anisole has also been shown to be produced by *Streptomyces albulus* and further tests showed pure anisole cause hyphal shriveling Wu et al. (44). Vacuolization was not observed in hyphae exposed to 2-undecanone, despite marked inhibition of fungal growth observed in our study as well as previous work on this bacteria volatile (45). Fungistatic effects may occur through multiple mechanisms and bacteria may be producing compounds with multiple modes of action. The overlap of several produced compounds by the different bacteria species supports this hypothesis; 2-undecanone was produced by all species, (although in lower amounts by both of the *Serratia* species). Anisole was produced by only *S. marcescens*, and dimethyl trisulfide was produced by both *Serratia* species, but at much higher amounts by *S. proteamaculans*.

Interestingly, we can see that the response of *R. stolonifer* to different bacterial species’ volatile profiles mirrors results of tests that expose fungi to headspace volatiles emanating from pure compounds. In the donut plate assays at 24 and especially 48 hours *B. subtilis* volatile inhibition capacity is lower than the other two *Serratia* species. In the pure volatile assays, we can see that the primary product of *B. subtilis*, 2-undecanone, slows but cannot prevent growth, unlike anisole and dimethyl trisulfide.

It is important to note that our study was carried out using bacteria growing on rich media, which has been optimized to support high bacteria population density. In the circular “donut plate” assays, biological activity of bacterial volatiles was only evident after 24 hours of bacteria growth, which indicates that a high quantity of cells is required to produce sufflcient volatiles to inhibit fungi. Likewise, the fungi were plated on optimal growth medium in the physically separated but common headspace. Thus our assays are only documenting fungistatic effects of bacterial volatiles under the most ideal conditions for both members of this microbe-microbe interaction. Even the *R. stolonifer* assays that used cut strawberries are rather idealized, as the strawberry substrate is suitable for the fast growing fungal colonizer. We feel these experiments still provide evidence that fungistatic activity can occur semi-natural conditions.

Testing pure compounds on *R. stolonifer* showed that not all compounds can inhibit even at much higher concentrations than detected from bacterial culture samples. For example, we recovered 5 - 10 ng of tropone (2,4,6-cycloheptatrien-1-one) from many samples, but even at 1000 times the concentration no significant inhibition was detected. Other compounds like dimethyl trisulfide and 2-undecanone have been shown to have minimal inhibitory concentrations (24 mg and 15 mg respectively) around the same levels we tested (10 mg). Further research is needed to document natural volatile emissions and concentrations, and measure fungal inhibition under realistic bacteria populations sizes in environments such as the soil in the rhizosphere, plant roots, or during co-colonization of rotten fruits.

These results provide useful insights into the diversity and potential functions of bacterial volatiles. For example, during headspace sampling that lasted only a few hours, we collected over a microgram of volatiles from many of our bacterial cultures. Thus, when sufficient resources are available, volatile production seems to be prioritized metabolically. This suggests that distance-based inhibition of competitors, including fungi, may be an important but somewhat overlooked element of microbial community assembly. Furthermore, most of the species used in our experiments produced more than one compound with fungistatic activity, which may increase the range of species against which volatile combinations are active or help to prevent the rapid evolution of resistance mechanisms. Although the identified compounds may not be the only fungistatic compounds in the volatile profile, they may prove useful for development of molecular genetic studies to understand the mechanisms of volatile production in bacteria. Future efforts should employ molecular and genetic tools to elucidate the biosynthetic production pathways of these compounds as well as develop an understanding of what genetic and environmental conditions influence volatile production. The outcomes of such explorations have the potential to lead to additional resources for fungal biocontrol in agriculture and postharvest crop protection, providing additional compounds which may have an impact on strategies for treating human and animal mycoses, and uncovering mechanisms of microbial competition and community assembly.

## SUPPLEMENTAL DATA

The following are the included supplemental data:

Supplemental Table 1 - Optical Density Growth Values
Supplemental Table 2 - ANOVA Posthoc Values
Supplemental Table 3 - Volatile Compound Measurements
Supplemental Figure 1 and Table 4 - https://doi.org/10.5281/zenodo.3926700

## ACKNOWLEDGMENTS

J.E.S., J.C., S.M., and D.C-H, were supported by National Science Foundation grant DEB-1441715 and United States Department of Agriculture - National Institute of Food and Agriculture Hatch project CA-R-PPA-5062-H to J.E.S. J.E.S. is a CIFAR fellow in the Fungal Kingdom: Threats and Opportunities program. K.E.M. was supported by Hatch project funds (CA-R-ENT-5144-H). The funders had no role in study design, data collection and interpretation, or the decision to submit the work for publication. The authors declare no conflicts of interest. We would like to thank Earl Kang for his help in designing the “donut” plate assay.

